# *Artemisia annua* L. extracts inhibit the *in vitro* replication of SARS-CoV-2 and two of its variants

**DOI:** 10.1101/2021.01.08.425825

**Authors:** M.S. Nair, Y. Huang, D.A. Fidock, S.J. Polyak, J. Wagoner, M.J. Towler, P.J. Weathers

## Abstract

**Ethnopharmacological relevance:** *Artemisia annua* L. has been used for millennia in Southeast Asia to treat “fever”. Many infectious microbial and viral diseases have been shown to respond to *A. annua* and communities around the world use the plant as a medicinal tea, especially for treating malaria.

**Aim of the Study:** SARS-CoV-2 (the cause of Covid-19) globally has infected and killed millions of people. Because of the broad-spectrum antiviral activity of artemisinin that includes blockade of SARS-CoV-1, we queried whether *A. annua* suppressed SARS-CoV-2.

**Materials and Methods:** Using Vero E6 and Calu-3 cells, we measured anti viral activity SARS-CoV-2 activity against fully infectious virusof dried leaf extracts of seven cultivars of *A. annua* sourced from four continents. IC_50_s were calculated and defined as (the concentrations that inhibited viral replication by 50%.) and CC50s (the concentrations that kill 50% of cells) were calculated.

**Results:** Hot-water leaf extracts based on artemisinin, total flavonoids, or dry leaf mass showed antiviral activity with IC_50_ values of 0.1-8.7 μM, 0.01-0.14 μg, and 23.4-57.4 μg, respectively. Antiviral efficacy did not correlate with artemisinin or total flavonoid contents of the extracts. One dried leaf sample was >12 years old, yet the hot-water extract was still found to be active. The UK and South African variants, B1.1.7 and B1.351, were similarly inhibited. While all hot water extracts were effective, concentrations of artemisinin and total flavonoids varied by nearly 100-fold in the extracts. Artemisinin alone showed an estimated IC_50_ of about 70 μM, and the clinically used artemisinin derivatives artesunate, artemether, and dihydroartemisinin were ineffective or cytotoxic at elevated micromolar concentrations. In contrast, the antimalarial drug amodiaquine had an IC_50_ = 5.8 μM. Extracts had minimal effects on infection of Vero E6 or Calu-3 cells by a reporter virus pseudotyped by the SARS-CoV-2 spike protein. There was no cytotoxicity within an order of magnitude above the antiviral IC_90_ values.

**Conclusions:** *A. annua* extracts inhibit SARS-CoV-2 infection, and the active component(s) in the extracts is likely something besides artemisinin or a combination of components that block virus infection at a step downstream of virus entry. Further studies will determine in vivo efficacy to assess whether *A. annua* might provide a cost-effective therapeutic to treat SARS-CoV-2 infections.

**List of compounds studied:**

Amodiaquine
Artemisinin
Artesunate
Artemether
Deoxyartemisinin
Dihydroartemisinin

**Highlights:**

- *Artemisia annua* is effective in stopping replication of SARS-CoV-2 including 2 new variants.
- The anti-viral effect does not correlate to artemisinin, nor to the total flavonoid content.
- The anti-viral mechanism does not appear to involve blockade virus entry into cell.
- The plant offers two additional benefits: a decreased inflammatory response and blunting of fibrosis.
- *A. annua* may provide a safe, low-cost alternative for treating patients infected with SARS-CoV-2.

## 1.0 INTRODUCTION

The global pandemic of SARS-CoV-2 (the etiologic agent of COVID-19) has infected over 110 million people and killed nearly 2.5 million as of February 19, 2021 (https://coronavirus.jhu.edu/). There is an intense effort to distribute the registered Pfizer/BioNTech, Moderna, and J&J vaccines, but to our knowledge, besides Remdesivir there is no approved, orally deliverable, small-molecule therapeutic and global infections keep rising with and new variants continue to emerge.

The medicinal plant *Artemisia annua* L. produces the antimalarial therapeutic artemisinin, a sesquiterpene lactone produced and stored in the glandular trichomes located on the shoots and particularly the leaves and flowers of the plant. Both the plant and artemisinin have been used safely for over 2,000 years to treat a variety of fever-related ailments, especially malaria (Hsu, 2006). Artemisinin derivatives (Figure 1) are front-line therapeutics for treating malaria and are delivered with a second antimalarial drug, such as lumefantrine or amodiaquine, which are formulated as artemisinin-based combination therapies (Blasco et al. 2017). Artemisinins also possess antiviral activity (Efferth 2018). Extracts of *A. annua* showed anti-SARS-CoV-1 activity, suggesting that they may be active against SARS-CoV-2 (Li et al. 2005).

**Figure 1.**
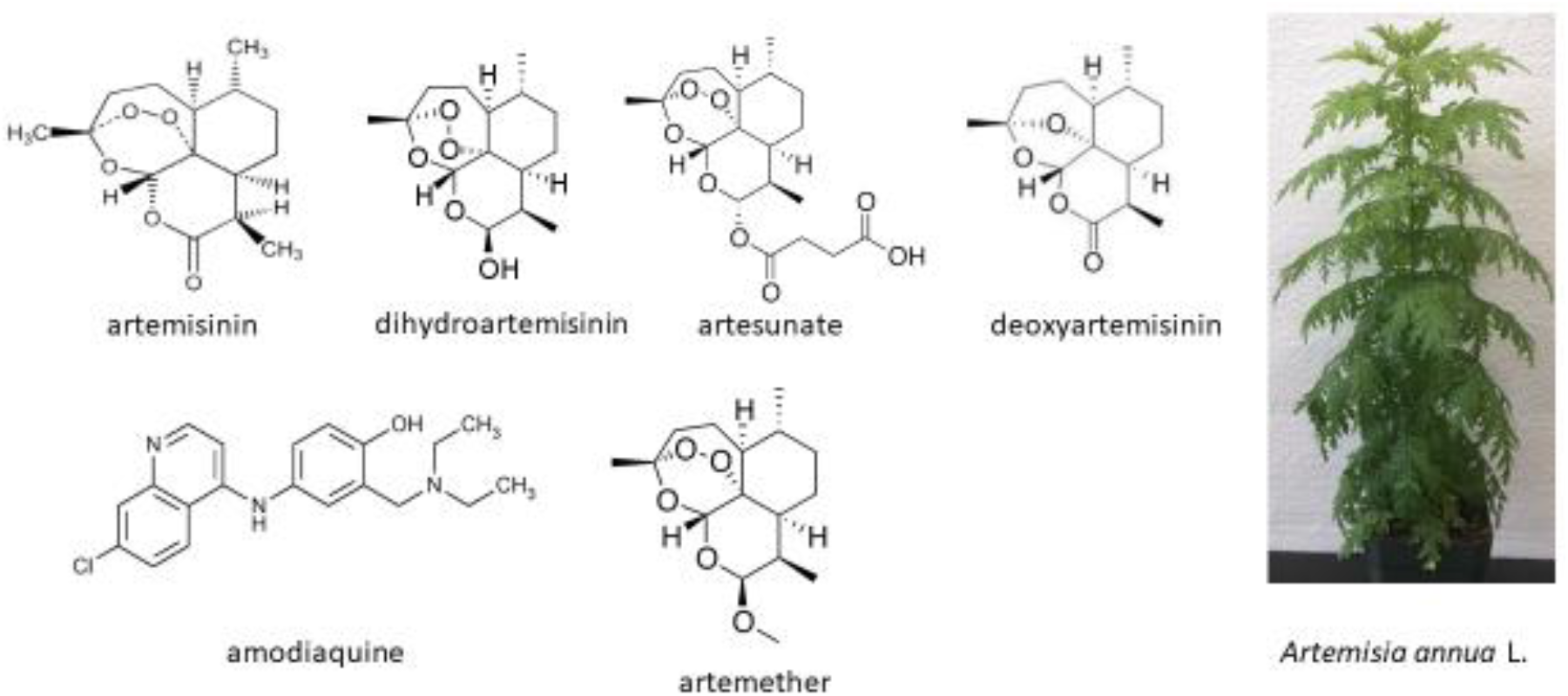
Compounds used in this study and the plant *Artemisia annua* L.

Artemisinin delivered directly from the consumption of *A. annua* leaf powder is highly bioavailable and distributes through peripheral blood and into a plethora of organs including lungs, liver, heart, and brain (Desrosiers et al. 2020). Furthermore, both artemisinins and the plant *A. annua* reduce levels of inflammatory cytokines including IL-6 and TNF-α *in vivo* (Desrosiers et al. 2020; Hunt et al. 2015; Shi et al. 2015). These effector molecules can be problematic during the “cytokine storm” suffered by many SARS-CoV-2 patients (Schett et al. 2020). Artemisinin also blunts fibrosis (Larson et al. 2019; Dolivo et al. 2020), another problem experienced by SARS-CoV-2 survivors that causes more lasting damage to organs (Lechowicz et al. 2020; Liu et al. 2020a). A recent report showed that a number of artemisinin-related compounds have some anti-SARS-CoV-2 activity, with dihydroartemisinin, artesunate, and arteannuin B having IC_50_ values <30 μM (Cao et al. 2020), and dihydroartemisinin ACTs having 1-10 μM IC_50_ values (Bae et al. 2020). Artesunate was reported to have IC_50_ values against SARS-CoV-2 of 7-12 μg/mL (0.7-1.2 μM; Gilmore et al. 2020) and 2.6 μM (Bae et al. 2020).

In a recent small human trial, Li et al. (2021) showed that artemisinin-piperaquine was safe and twice as effective as placebo in completely eliminating the virus 21 days after treatment for seven days. Knowing that artemisinin is much more bioavailable *per os* when delivered via *A. annua* (Weathers et al. 2011; Weathers et al. 2014; Desrosiers et al. 2020), we posited that encapsulated powdered dried leaves of *A. annua* may be a safe, cost-effective therapeutic to combat SARS-CoV-2 infections.

Here we report *in vitro* results from testing extracts of a diversity of *A. annua* cultivars against infection of Vero E6 and Calu-3 cells by fully infectious SARS-CoV-2 and two of its recent variants, with correlation analyses of antiviral IC_50_ efficacy to artemisinin and total flavonoid contents.

## 2.0 METHODS

### 2.1 Plant material, extract preparations, and artemisinin and total flavonoid analyses

Batches of dried leaves of various cultivars of *Artemisia annua* L. with source, age, and voucher identity when known are shown in Table 1. Hot-water extracts (tea infusions) were prepared as follows: dried leaves at 10 g/L were added to boiling water on a stir plate and boiled for 10 min, then poured through a 2 mm stainless steel sieve to retain most solids. Extracts were then cooled and sterile-filtered (0.22 μm) prior to being stored at −20°C. Dichloromethane (DCM) extracts of dried leaves were also prepared by extraction of 25 mg in 4 mL DCM for 30 min in a sonicating water bath (Fischer Scientific FS60, 130 W), separating solvent from solids with Pasteur pipets, drying under nitrogen flow, and storing at −20°C until analyzing for artemisinin using gas chromatography / mass spectrometry, as detailed in Martini et al. (2020). For artemisinin analysis of tea infusions, two-phase overnight aliquots extracted in DCM in a 1:1 ratio were separated by using Pasteur pipets, dried under nitrogen flow, and stored at −20°C until analysis as previously noted (Martini et al. 2020). Total flavonoids were analyzed in DCM extracts via the aluminum chloride method of Arvouet-Grand et al. (1994) and were quantified as quercetin equivalents. Artemisinin and total flavonoid contents of tea infusions are shown in Table 2. The DCM extract of *A. annua* (cv. SAM) contained a total of 34 mg of artemisinin. After solubilizing in PEG400 containing 5% DMSO the concentration was 8.95 mg/mL.

**Table 1.**
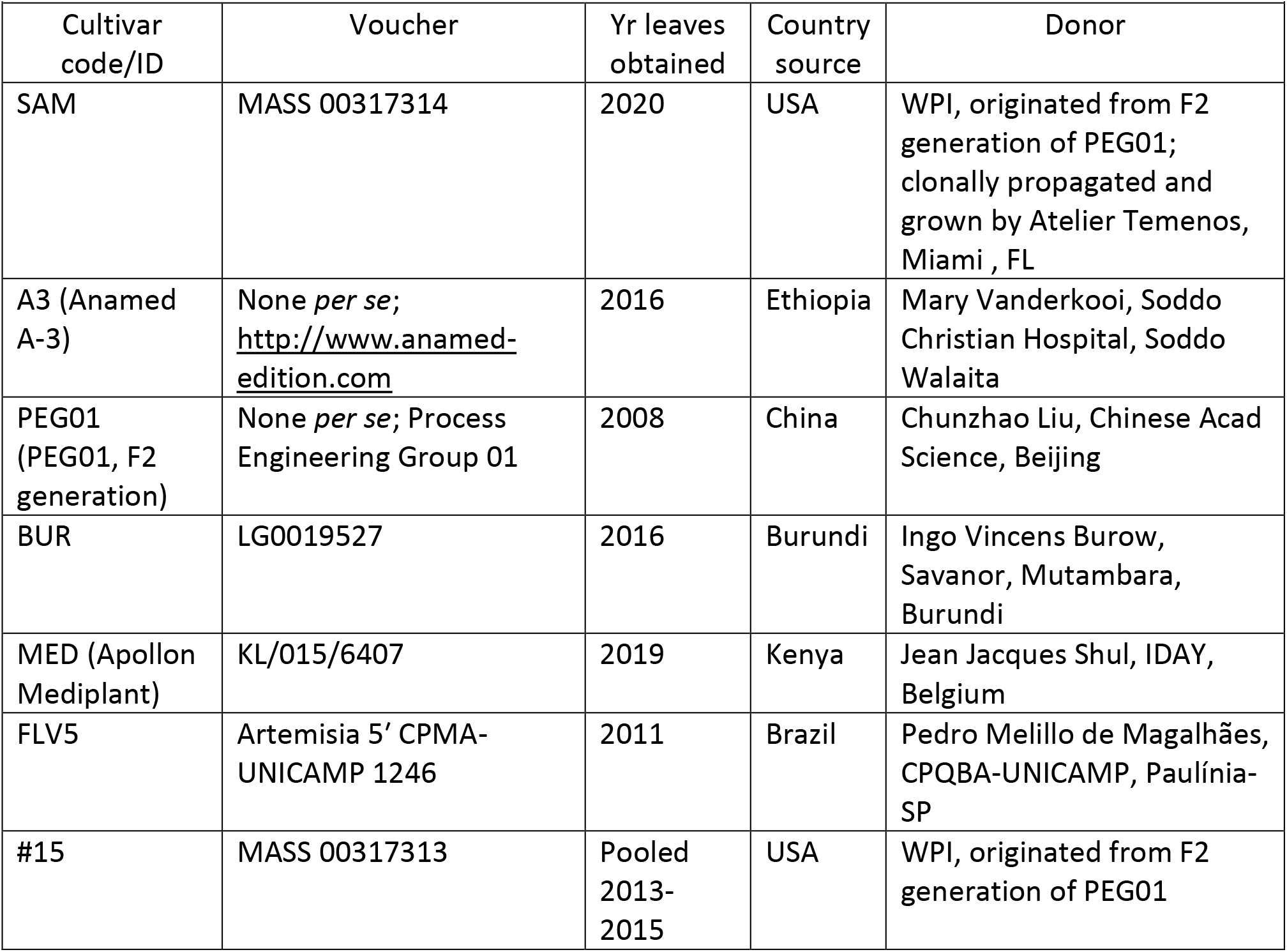
*Artemisia annua* L. cultivars used in the SARS-CoV-2 antiviral analyses.

|  |  |  |  |  |
| --- | --- | --- | --- | --- |
| Cultivar code/ID | Voucher | Yr leaves obtained | Country source | Donor |
| SAM | MASS 00317314 | 2020 | USA | WPI, originated from F2 generation of PEG01; clonally propagated and grown by Atelier Temenos, Miami , FL |
| A3 (Anamed A-3) | None *per se*; http://www.anamed-edition.com | 2016 | Ethiopia | Mary Vanderkooi, Soddo Christian Hospital, Soddo Walaita |
| PEG01 (PEG01, F2 generation) | None *per se*; Process Engineering Group 01 | 2008 | China | Chunzhao Liu, Chinese Acad Science, Beijing |
| BUR | LG0019527 | 2016 | Burundi | Ingo Vincens Burow, Savanor, Mutambara, Burundi |
| MED (Apollon Mediplant) | KL/015/6407 | 2019 | Kenya | Jean Jacques Shul, IDAY, Belgium |
| FLV5 | Artemisia 5' CPMA-UNICAMP 1246 | 2011 | Brazil | Pedro Melillo de Magalhães, CPQBA-UNICAMP, Paulínia-SP |
| #15 | MASS 00317313 | Pooled 2013-2015 | USA | WPI, originated from F2 generation of PEG01 |

**Table 2.** Calculated IC_50_ and IC_90_ values for the 7 *A. annua* cultivars against Vero E6 cells infected with SARS-CoV-2 USA/WA1 (MOI 0.1) based on their measured artemisinin (ART), total flavonoids (tFLV), and dry leaf mass content (DW) in the hot water extracts.

|  |  |  |  |  |  |  |  |  |  |
| --- | --- | --- | --- | --- | --- | --- | --- | --- | --- |
| Sample ID | Artemisinin |  |  | Total Flavonoids |  |  | Dry *A. annua* Leaf Mass |  |  |
|  | ART in tea (µg/mL ± SE) | IC<sub>50</sub> (µM ART) | IC<sub>90</sub> (µM ART) | tFLV** (µg/mL ± SE) | IC<sub>50</sub> (µg) | IC<sub>90</sub> (µg) | Leaves extracted (g/L) | IC<sub>50</sub> (µg DW) | IC<sub>90</sub> (µg DW) |
| SAM1* (-20C) | 149.4 ± 4.4a | 8.7 | 18.8 | 35.4 ± 0.2a | 0.13 | 0.28 | 10 | 34.9 | 75.2 |
| SAM2* (4C) | 131.6 ± 3.4b | 5.9 | 12.3 | 37.2 ± 0.7a | 0.14 | 0.29 | 10 | 38.4 | 79.0 |
| A3 | 42.5 ± 1.8c | 1.4 | 2.7 | 10.5 ± 0.3b | 0.03 | 0.06 | 10 | 28.9 | 54.9 |
| PEG01 | 82.7 ± 2.8d | 3.2 | 13.6 | 17.6 ± 0.6c | 0.06 | 0.25 | 10 | 32.9 | 139.3 |
| FLV5 | 73.3 ± 2.5d,e | 4.9 | 14.5 | 7.9 ± 0.1b | 0.07 | 0.21 | 10 | 57.4 | 167.8 |
| #15 | 47.8 ± 2.5c,f | 1.8 | 5.4 | 10.7 ± 0.2b | 0.05 | 0.15 | 10 | 32.3 | 95.7 |
| BUR | 20.1 ± 0.8g | 0.1 | 0.2 | 7.3 ± 0.2b | 0.01 | 0.03 | 10 | 13.5 | 37.7 |
| MED | 59.4 ± 1.6e,f | 0.4 | 1.1 | 22.3 ± 0.5d | 0.05 | 0.13 | 10 | 23.4 | 58.7 |

\* SAM1 and SAM2 are replicated hot water extracts from the same batch of *A. annua* leaves grown and processed from Atelier Temenos; SAM1 was stored at -20C, thawed and reanalyzed at the same time as SAM2. Data are the average of ≥ 6 independently extracted leaf samples. \*\*
Quercetin equivalents.
Letters in ART and tFLV columns indicate statistical significance at $P \leq 0.05$ .

### 2.2 Viral culture and analyses

Vero E6 cells, obtained from the American Type Culture collection (ATCC CRL-1586), were cultured in Essential Minimal Eagle’s Medium (EMEM) containing penicillin-streptomycin (1x 100 U/mL) and 10% fetal calf serum. SARS-CoV-2 isolate USA/WA12020, UK variant B1.1.7 (CA_CDC_5574/2020), and South African variant B1.351 (hCoV-19/South Africa/KRISP-EC-K005321/2020) were from BEI Resources (www.beiresources.org). We infected Vero E6 cells with the USA/WA1 isolate according to Liu et al. (2020b). Briefly, infected cells were incubated in flasks until a viral cytopathic effect was observed. The supernatant was then harvested and titered for its tissue culture infective dose (TCID) using an end point dilution method. TCID was calculated using the Reed-Muench proportional distance method (Reed and Muench 1938). Viral aliquots were frozen, then later thawed and used for infection experiments at their desired infectivity (multiplicity of infection (MOI).

### 2.3 Assays for determining drug inhibition of SARS-CoV-2

Except for tea infusions that were diluted in water and used directly, amodiaquine, artesunate, artemether, artemisinin, deoxyartemisinin, and dihydroartemisinin compounds were solubilized and diluted in 5% DMSO in PEG400 or 5% DMSO in EMEM enriched with fetal calf serum at a final concentration of 7.5%, prior to testing for efficacy against SARS-CoV-2. Indicated dilutions of the drug were incubated for 1 h in wells of 96 well tissue culture plates containing a monolayer of Vero E6 cells seeded the day before at 20,000 cells/well. Post incubation of the drug with the cells, SARS-CoV-2 USA/WA1 virus was added to each well at a multiplicity of infection of 0.1. Cells were cultured for 3 days at 37°C in 5% CO_2_ and scored for cytopathic effects as detailed in Liu et al. (2020b). Vesicular Stomatitis Virus (VSV)-spike pseudoviruses were generated as described (Hoffmann et al. 2020; Whitt 2010), using the spike gene from SARS-CoV-2 containing the D614G mutation (Korber et al. 2020). The construct also contains a deletion of 18 amino acids from the C-terminus, which facilitates loading onto pseudovirus particles. The construct (Δ18 D614G) was kindly provided by Markus Hoffmann and Stefan Pöhlmann (Leibniz-Institut für Primatenforschung, Germany). The day prior to infection, Vero E6 and Calu-3 cells (ATCC HTB-55) were plated in black, clear-bottomed plates at 10,000 and 30,000 cells/well, respectively, in a final volume of 90 μl. Cells were then treated with 10 μl of serially diluted *Artemisia* extract in water and incubated for 1 h prior to infection with 100 μl of VSV-spike Δ18 D614G pseudovirus. At 22 h post-infection, PrestoBlue was added 2 h before the end of assay, so that cell viability in parallel non-infected, drug-treated wells could be measured. Virus-produced Renilla luciferase activity was measured by Renilla-Glo assay at 24 h post-infection. Results were converted into percent of control. Drug concentrations were log transformed and the concentration of drug(s) that inhibited virus by 50% (*i.e.*, IC_50_), and the concentration of drug(s) that killed 50% of cells (*i.e.*, CC_50_), were determined via nonlinear logistic regressions of log(inhibitor) versus response-variable dose-response functions (four parameters) constrained to a zero-bottom asymptote by statistical analysis using GraphPad Prism 9 (GraphPad Software, Inc.) as described by Hulseberg et al. (2019).

### 2.4 Cell viability assay

To determine the viability of Vero E6 cells post drug treatment, cells were exposed to indicated doses of tea infusions diluted in EMEM containing fetal calf serum at a final concentration of 7.5% and incubated at 37°C in 5% CO_2_ for 24 h. Cells were then washed and treated with 100 μL XTT reagent premixed with activation agent, followed by incubation for another 2 h at 37°C in 5% CO_2_. Culture medium was removed, and absorbance measured at 450 nm. The absorbance ratio of treated to untreated cells was plotted as percent viability. Imatinib, an FDA-approved apoptosis inducer and tyrosine kinase inhibitor, was used as a positive control.

### 2.5 Chemicals and reagents

Unless otherwise stated all reagents were from Sigma-Aldrich (St. Louis, MO). DCM was from ThermoFisher (Waltham, MA, USA); artemisinin was from Cayman Chemical (Ann Arbor, MI, USA); artemether, artesunate, and dihydroartemisinin were gifts from Prof. J. Plaizier-Vercammen (Brussels, Belgium); deoxyartemisinin was from Toronto Research Chemicals (North York, ON, Canada); amodiaquine HCl hydrate (Cat #: 562290) and imanitib (Cat # 100956) were from Medkoo Biosciences Inc. (Morrisville, NC, USA); EMEM (Cat # 30-2003) and XTT reagent (Cat # 30-1011k) were from ATCC; PrestoBlue was from Life Technologies (Cat #P50201); Renilla-Glo was from Promega (E2720).

### 2.6 Statistical analyses

All *in vitro* anti-SARS-CoV-2 analyses were done at least in triplicate. Plant extract analyses had n≥6 independent assays. IC_50_ and IC_90_ values were calculated using GraphPad Prism V8.0 or V9. Correlations between antiviral activity and artemisinin or total flavonoids used Spearman’s Rho analysis (Spearman 1904). Statistical significance of artemisinin and total flavonoid content in hot water extracts was calculated via ANOVA using GraphPad Prism V8.0.2.

## 3.0 RESULTS

### 3.1 *Artemisia annua* hot water extracts have anti-SARS-CoV-2 activity

Hot water extracts of the *A. annua* cultivars used in the study had significantly different artemisinin contents ranging from 20.1 ± 0.8 to 149.4 ± 4.4 μg/mL (Table 2). Total flavonoid content of leaf material ranged from 7.3 ± 0.2 to 37.2 ± 0.7 μg/mL (Table 2). All cultivars showed anti-SARS-CoV-2 activity (Figure 2; Table 2), and IC_50_ values calculated on the basis of artemisinin or total flavonoid content ranged from 0.1-8.7 μM, or 0.01-0.14 μg/mL, respectively (Table 2). On the basis of leaf dry mass, IC_50_ values ranged from 13.5-57.4 μg dry weight (DW). On a μg artemisinin/mL tea basis, the IC_50_ of the samples ranged from 0.03 to 2.5 μg/mL. Analysis of frozen (SAM −20C) extracts remained potent upon thawing and reanalysis (Table 2, Figure 2). Leaf samples that were 12 years old were also active with an IC_50_ of 32.9 μg DW. Notably two recently isolated variants of SARS-CoV-2 from the UK (B1.1.7) and South Africa (B1.351) that are of concern due to the lowered effect of vaccines and antibodies against them (Wang et al. 2021) were similarly susceptible to *A. annua* extracts from BUR, MED, A3 and SAM1 (Figure 3) with IC_50_s and IC_90_s within the range of those values measured for the original isolate from the US (Table 2).

**Figure. 2.**
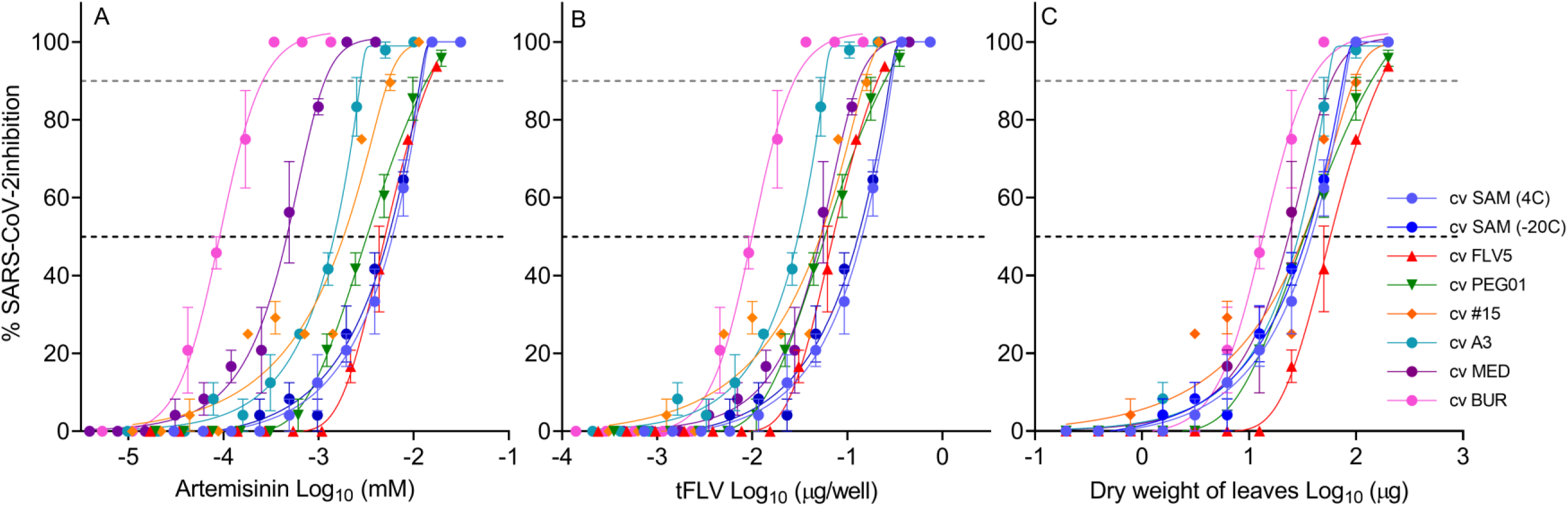
IC_50_ inhibition plots of extracts for efficacy against Vero E6 cells infected with SARS-CoV-2 USA/WA1 (MOI 0.1) based on: artemisinin (A); total flavonoids (tFLV) (B); or dry mass of *A. annua* leaves (C) used in the experiments. SAM −20C = SAM1; SAM 4C = SAM2. Data are plotted from an average of three replicates with ± SE.

**Figure 3.**
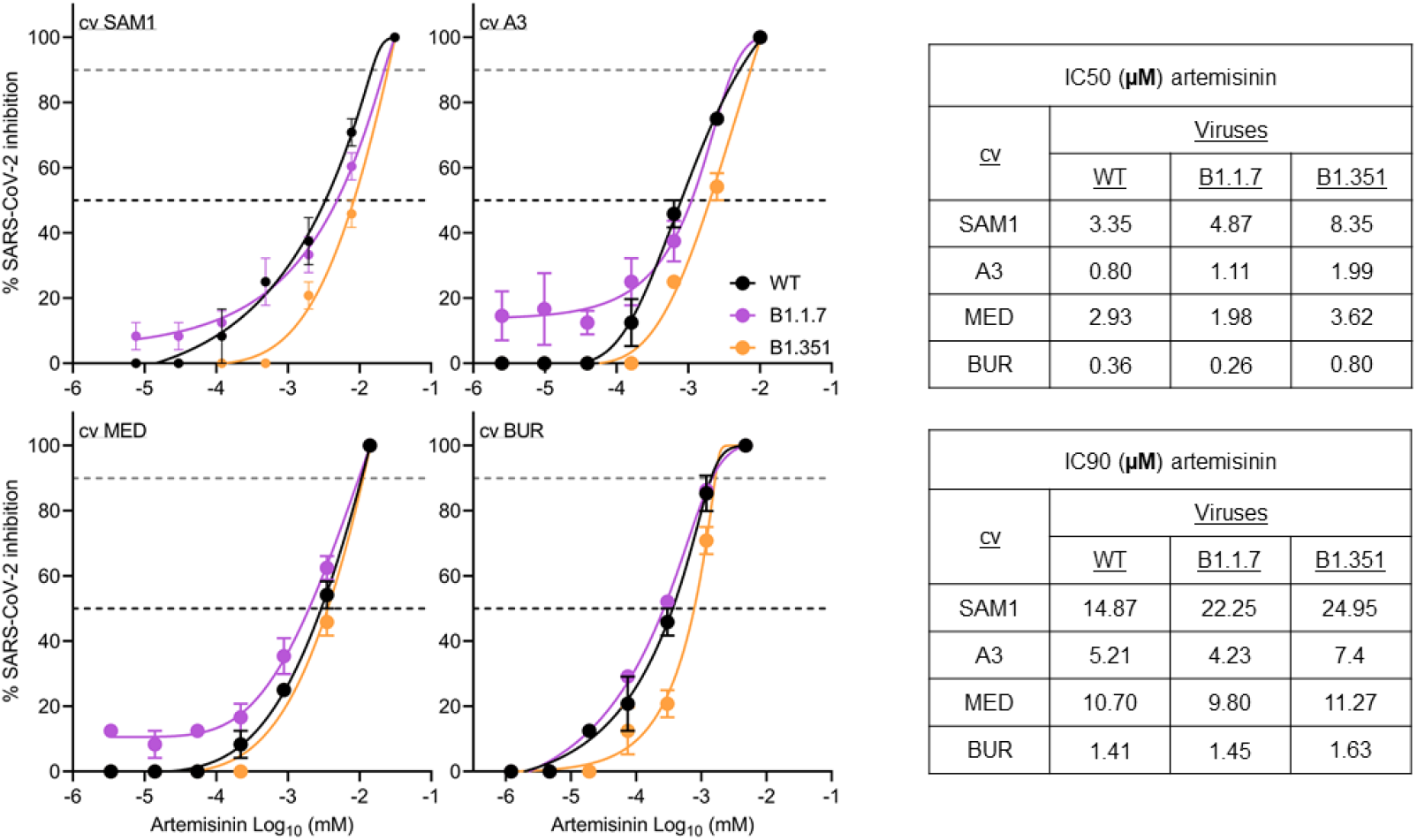
IC_50_ inhibition plots and IC_50_ and IC_90_ values for four *A. annua* cultivar extracts for efficacy against Vero E6 cells infected with WT (USA/WA12020) SARS-CoV-2 and variants, B1.1.7 and B1.351 (MOI 0.1) based on their measured artemisinin in the hot water extracts. Data are plotted from an average of three replicates with ± SE.

Infection of Vero E6 or Calu-3 human lung cells by VSV-spike pseudoviruses was minimally inhibited by the extract, except perhaps at the highest artemisinin dose tested of 500 μg/mL (Figure 4). Indeed, GraphPad Prism-calculated IC_50_/CC_50_ values were 545/3564 μg/mL for Calu-3 and 410/810 μg/mL for Vero E6 cells.

**Figure 4.**
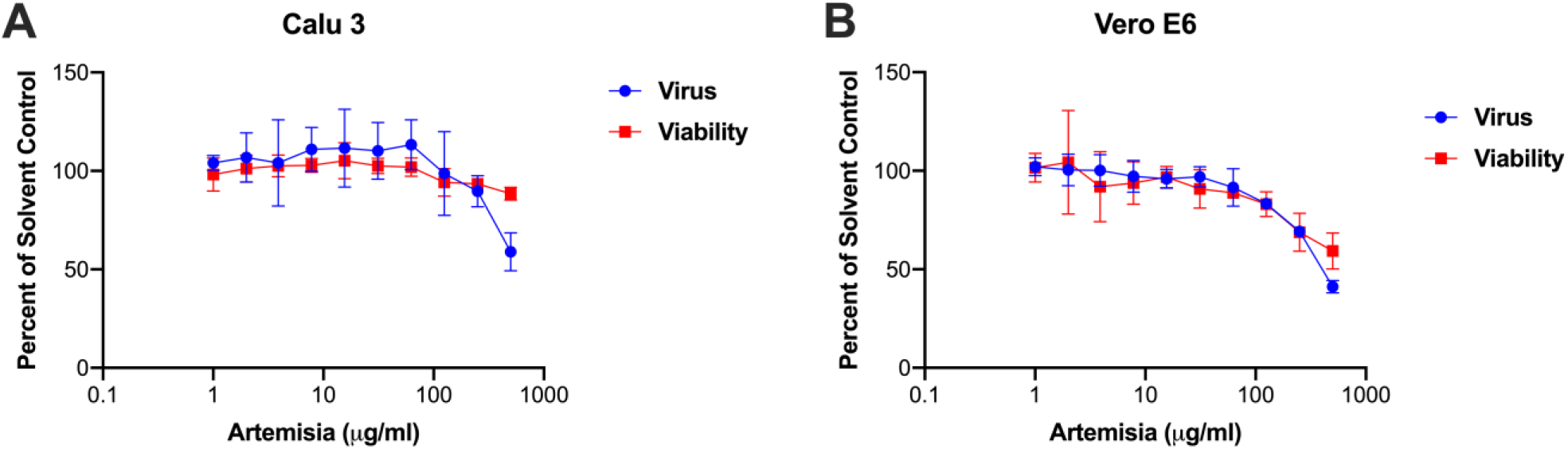
VSV spike pseudovirus in Calu-3 and Vero E6 cells and their viability in response to increasing hot water *Artemisia* extracts as percent of solvent controls. *Artemisia* concentration refers to dry leaf mass extracted with hot water. Data plotted using nonlinear regression curve fitting using GraphPad Prism. Data are averages of triplicate samples per condition and error bars are ± SD. Data are a representative experiment that was repeated twice.

### 3.2 Hot water extracts are not cytotoxic

When cytotoxicity of the hot water extracts to the Vero E6 and Calu 3 cells was measured, cell viability did not substantially decrease (Figures 4 and 5) at 24 h post treatment. In comparison, the apoptotic inducer imatinib showed a dose-dependent decrease in viability of the cells by 90% (Figure 5 inset). At the higher concentrations of hot water extracts, there appeared to be proliferation of Vero E6 cells (Figure 5).

**Figure 5.**
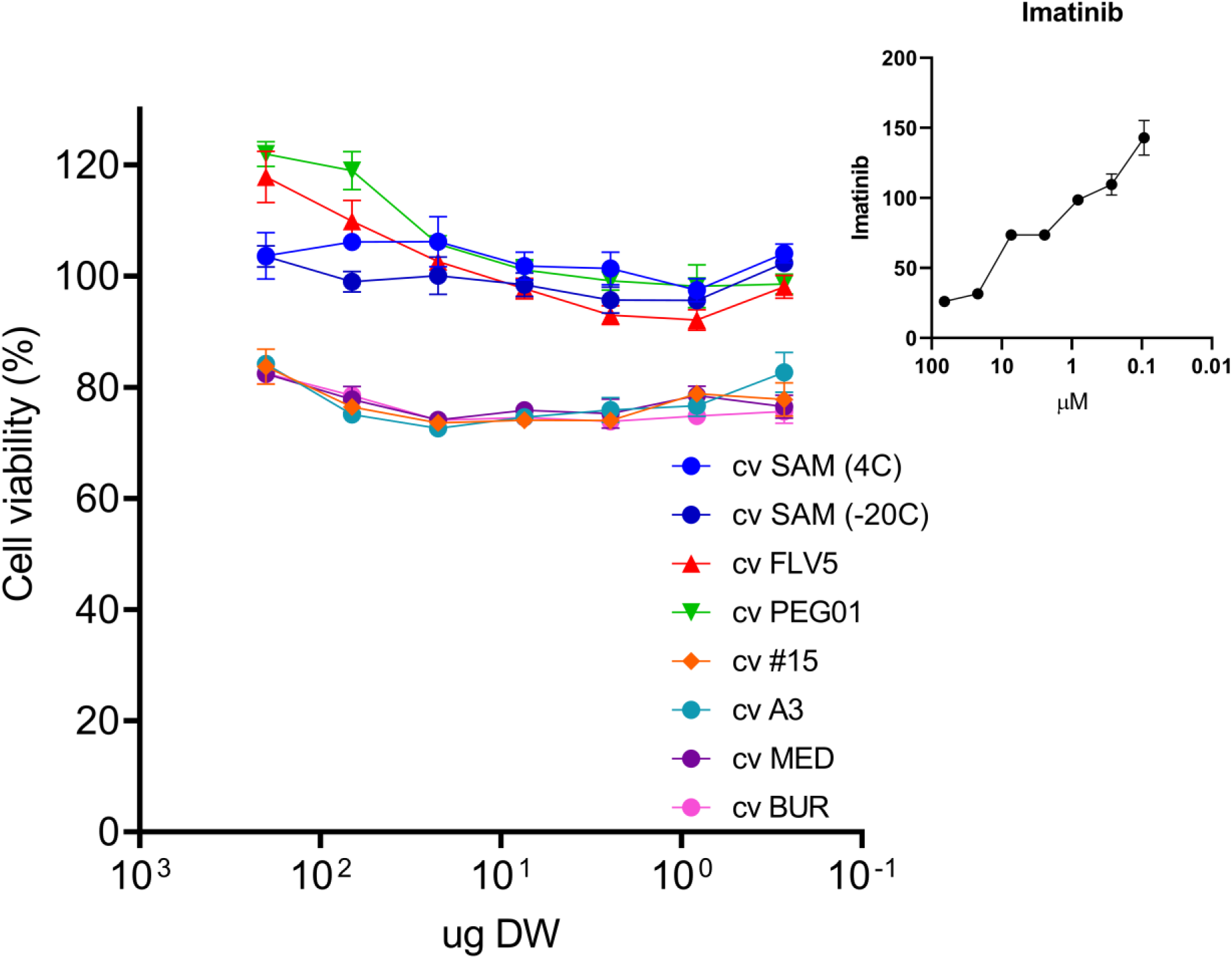
Cytotoxicity of Vero 6 cells in response to imatinib (A) and *A. annua* hot water extracts (B). Data are plotted from an average of three replicates with ± SE.

### 3.3 Activity of antimalarials

In a separate analysis, DCM and hot water extracts of *A. annua* were compared, yielding IC_50_ values of 12.0 and 11.8 μM, respectively (Figure 6). However, due to solvent toxicity at higher concentrations of the drug on Vero E6 cells, the IC_50_ of the DCM extract had to be estimated at 12 μM. Similar solvent toxicity was encountered with artemisinin that subsequently was estimated to have an IC_50_ of 70 μM (Figure 6). Artemether efficacy also had to be estimated at 1.23 μM and was cytotoxic at concentrations slightly above that level (Figure 6). Artesunate and dihydroartemisinin were inactive at all tested concentrations. In contrast, amodiaquine showed efficacy at 5.8 μM (Figure 6).

**Figure 6.**
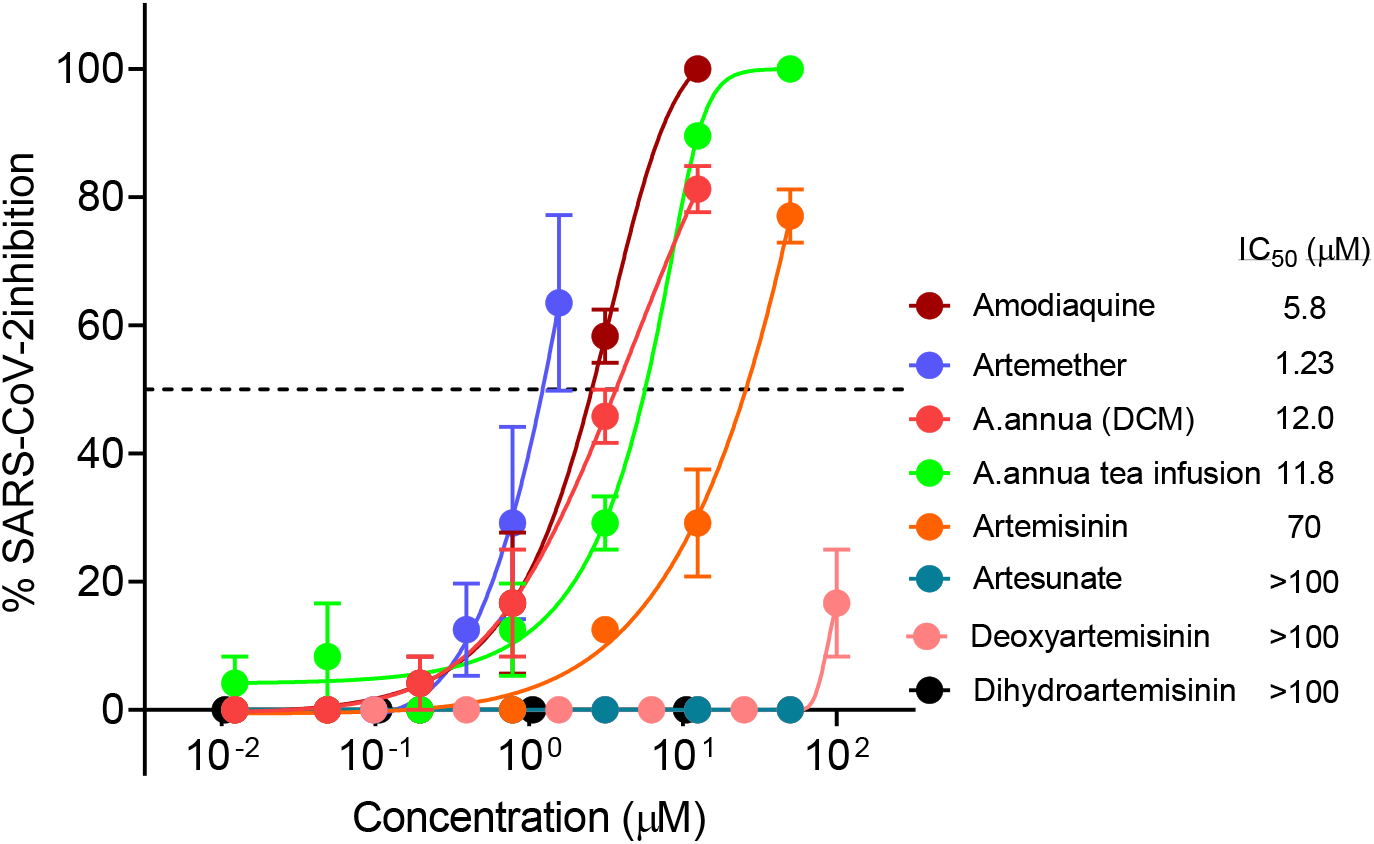
Comparison of *A. annua* SAM extracts and other antimalarial and artemisinin compounds against Vero E6 cells infected with SARS-CoV-2 USA/WA1 (MOI 0.1). A full concentration series for all samples except for the *A. annua* tea could not be fully tested due to solvent toxicity, which was also observed for *A. annua* in dichloromethane (DCM) at higher concentrations. Data are plotted from an average of three replicates with ± SE.

### 3.4 Anti-SARS-CoV-2 activity of hot water extracts inversely correlated to artemisinin or total flavonoid content

The IC_50_ quantifies the antiviral efficacy of a drug or extract. The lower the IC_50_, the more effective a drug or extract. To begin to define the bioactive components in *A. annua* responsible for suppressing SARS-CoV-2 infection, we correlated IC_50_ and IC_90_ (the concentration of drug that inhibits 90% of virus) with the artemisinin content of our extracts. A Spearman’s Rho analysis showed that both IC_50_ and IC_90_ values of the hot water extracts increased with with artemisinin and total flavonoid content (Figure 7). If artemisinin was the principle bioactive responsible for suppressing virus infection, then IC50 and IC90 concentrations should decrease with increasing concentrations of artemisinin, but they did not. Moreover, results of IC_50_ and IC_90_ calculations based on dry leaf mass used to prepare the tea were tightly grouped (Figure 2). Although cultivar IC_50_ ranking from most to least effective on dry weight basis was BUR, MED, A3, #15, PEG01, SAM1, SAM2, and FLV5 (Table 2), the maximum differential was less than 44 μg DW of dried leaves, or ~4.4 μL of tea infusion, an inconsequential difference. Collectively, the data suggest that artemisinin is not the principal molecule responsible for suppression of SARS-CoV-2 infection.

**Figure 7.**
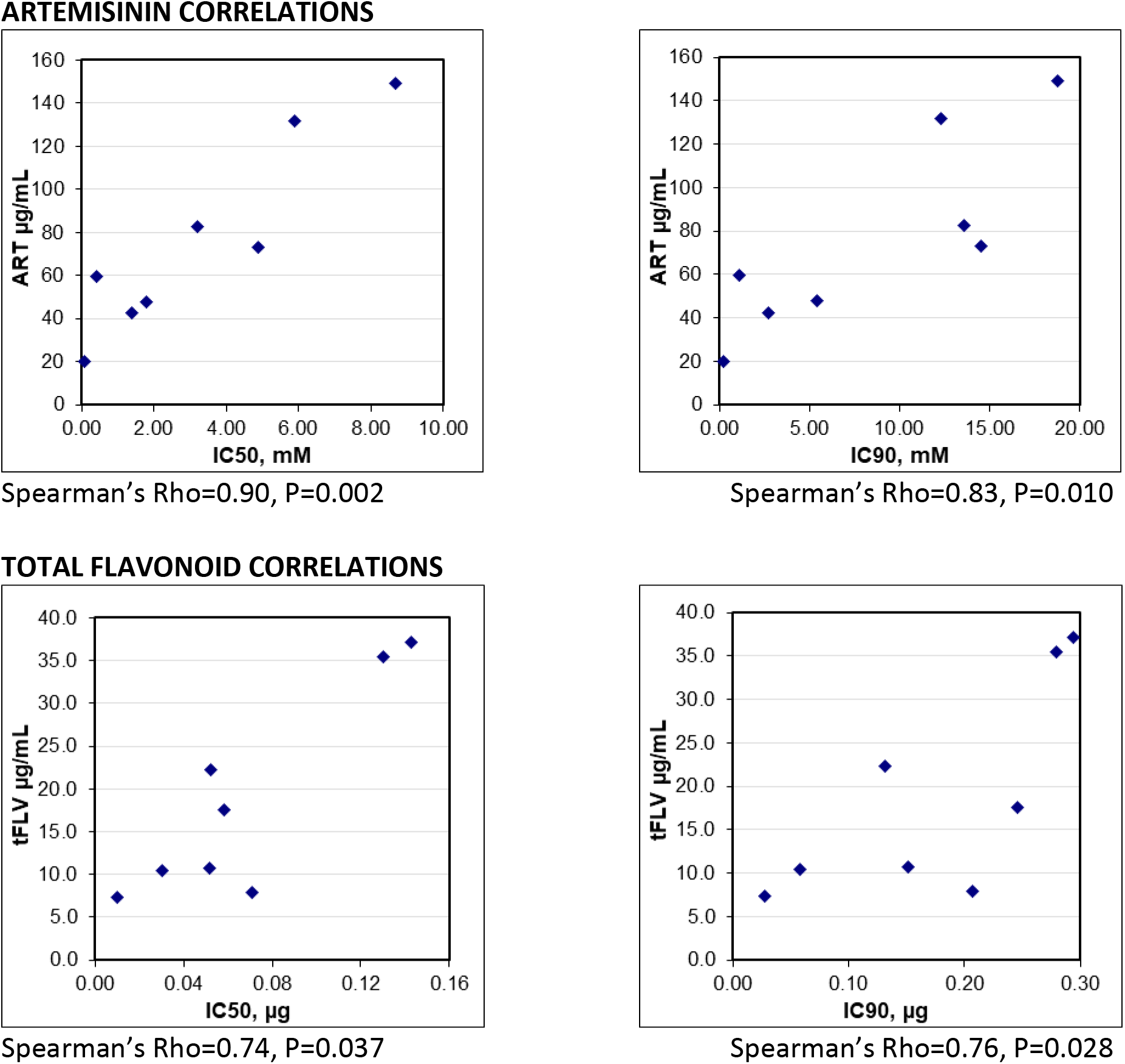
Spearman’s correlation scatter plots between artemisinin concentration or total flavonoid levels vs. calculated IC_50_ and IC_90_ for the hot water extract of each cultivar from data in Table 2.

### 3.5 Human bioavailability

For a preliminary query of the potential of using dried leaf *A. annua* (DLA) as a potential therapeutic, we tracked artemisinin as a marker molecule post consumption of *per os* delivered DLA to one human volunteer. One of us (PJW) consumed 3 g of encapsulated DLA of the SAM cultivar and had blood drawn at 2 and 5 h post consumption, resulting in serum measurements of 7.04 and 0.16 μg artemisinin/mL serum, respectively (See Supplemental Data).

Thus, at 2 h post ingestion, 36% of the original DLA-delivered artemisinin was detected in the serum, decreasing to 0.8% at 5 h post ingestion (See Supplemental Material for methodology and data Table S1). This corresponded at 2 h to 2.35 μg artemisinin/mL serum of DLA-delivered artemisinin per gram of DLA consumed.

## 4.0 DISCUSSION

This is the first report of anti-SARS-CoV-2 efficacy of hot water extracts of a wide variety of cultivars of *A. annua* sourced from four continents. These extracts had an IC_50_ corresponding to <12 μM artemisinin, with DCM extracts of *A. annua* showing similar efficacy. There was a similar response against the two variants when challenged by four of the extracts including three of the most efficacious. In contrast, artemisinin, when tested as a solo drug, had an estimated IC_50_ about sixfold greater (~70 μM), suggesting the plant extracts were more potent against SARS-CoV-2. Furthermore, the anti-SARS-CoV-2 effect was positively correlated to the artemisinin content of the extracts that varied by one to nearly two orders of magnitude for IC_50_ and IC_90_ values. A lower IC_50_ indicates a more active drug. Rather, the Spearman analysis showed that as the IC_50_ increased so also did the correlating artemisinin concentration. If artemisinin were the primary active component, one would expect the IC_50_, the concentration of drug that suppresses virus by 50%, to decrease with increasing concentrations of artemisinin, but the reverse occurred. Thus, our results suggest a possible antagonistic role of artemisinin in these extracts.

Total flavonoid content also similarly correlated to antiviral activity; IC_50_ and IC_90_ values increased with increasing flavonoid content. If total flavonoids were the primary active components, one would expect the IC_50_, and IC_90_ values to decrease with increasing concentrations of flavonoids, but they did not. One of the cultivar samples was obtained in 2008 and was still active at a level comparable to the most recently harvested cultivar samples, suggesting that the active principles are ubiquitous to different *A. annua* cultivars and chemically stable during long-term room temperature dry storage. None of the plant extracts were cytotoxic to Vero 6 or Calu-3 cells at concentrations approaching the IC_50_ or IC_90_ values. Indeed, at higher extract concentrations there was a slight increase in cell growth of Vero E6 cells, a positive response to *A. annua* extracts that also was reported for other healthy mammalian cells, e.g. fibroblasts (Rassias and Weathers 2019). Finally, the minimal antiviral effects against VSV pseudoviruses containing the SARS-CoV-2 spike protein suggests that *A. annua* inhibits SARS-CoV-2 infection primarily by targeting a post-entry step.

Although Cao et al. (2020) reported an EC_50_ of 10.28 μM for arteannuin B, a metabolite that is formed in a side branch of the artemisinin biosynthetic pathway and that is often present in *A. annua* extracts, only three of the tested tea extracts had any detectable arteannuin B with SAM having 3.2 μg/mL. Arteannuin B in BUR and MED was barely detectable. Thus, arteannuin B is tentatively eliminated as the principle active component, although if present in an extract, arteannuin B may be providing some antiviral effect as part of the more complex plant extract mixture. Although they can be present in substantial amounts in *A. annua* (Weathers and Towler 2014; Towler and Weathers 2015; see supplemental Table S2), neither artemisinic acid nor deoxyartemisinin, also metabolites in the artemisinin biosynthetic pathway, showed anti-SARS-CoV-2 activity in this study.

There is some discrepancy among IC_50_ molar values in this and other studies for anti-SARS-CoV-2 efficacy (Table 3). In contrast to Bae, Cao, and Gilmore, we did not observe any anti-SARS-CoV-2 activity for artesunate or dihydroartemisinin. Artemether in our study had an IC_50_ of 1.23 μM, while Cao et al. (2020) reported an EC_50_ of 73.8 μM but with less toxicity than we observed. In particular, we noted cytotoxicity of artemether. The contrasts are likely the result of differences in how we conducted our viral challenge experiments or solvents used to challenge the virus in Vero E6 cells. For example, our study solubilized our pure artemisinin and other antimalarial compounds in 5% DMSO in PEG400, while the other two studies solubilized compounds in DMSO. Our preliminary experiments indicated that solubilizing in pure DMSO was too toxic to Vero cells to achieve dosing of drug concentrations needed to obtain an IC_50_ value. In addition, Cao et al. also had a different viral assay system. We used an endpoint assay to measure the cytopathic effect of the replicating virus at 72 h and estimate the IC_50_ values while they collected supernatants to assay the total RNA levels at 24 h post infection using RT-PCR. We recognize that such inherent variations in the biological assays would offset the calculated values. Gilmore et al. (2020) also tested a hot water extract of *A. annua* and observed EC_50_ values ranging from 260-390 μg extract/mL. None of these studies reported IC_50_s based on the artemisinin content of their extracts.

**Table 3.** Comparative IC/EC_50_s for artemisinin derivatives and partner drug antimalarials.

Quercetin equivalents. Letters in ART and tFLV columns indicate statistical significance at $P \leq 0.05$ .
|  |  |  |  |  |  |
| --- | --- | --- | --- | --- | --- |
| Compound | Cao et al.<br>(2020) | Gilmore et al.<br>(2020) | Bae et al.<br>(2020) | Weston et<br>al. (2020) | This report |
| | $\mu\text{M}$ | | | | |
| Artemisinin | 64.5 | 534.8 | NM | NM | 70 |
| Arteannuin B | 10.3 | NM | NM | NM | NM |
| Artemisinic acid | >100 | NM | NM | NM | NM |
| Deoxyartemisinin | NM | NM | NM | NM | >100 |
| Dihydroartemisinin | 13.3 | NM | NM | NM | >100 |
| Artesunate | 13.0 | 18.2 | 53, 1.8<br>(Vero E6,<br>Calu-3) | NM | >100 |
| Arteether | 31.9 | NM | NM | NM | NM |
| Artemether | 73.8 | >600 (VeroE6)<br>453 (Huh7.5) | NM | NM | 1.2 |
| Artemisone | 49.6 | NM | NM | NM | NM |
| Amodiaquine | NM | NM | NM | 2.6-5.6 | 5.8 |
| Lumefantrine | 23.2 | NM | NM | NM | NM |

NM = not measured

Our hot water extracts are not directly comparable to those of Gilmore et al. because we did not dry, concentrate, and then weigh our extracts. Furthermore, we extracted for 10 min in boiling water, while they extracted for 200 min in boiling water. At present, it is not possible to compare our hot water extracts directly. In addition, different viruses were used in our study versus that of Gilmore et al., which could affect the inherent replication kinetics of the assay and in turn affect the specific IC_50_ numbers. In Nie et al. (2021), the same team tested hot water extracts prepared for a shorter time, but they still concentrated their extracts before testing. In contrast, we tested our tea extracts directly without any concentration. Interestingly, Nie et al. also reported efficacy of *Artemisia afra*, a perennial native to South Africa that produces no artemisinin (du Toit and van der Kooy, 2019). Extracts of that species were more effective (EC_50_ = 0.65 mg/mL) than pure artemisinin (EC_50_ =4.23 mg/mL) and extracts of four *A. annua* cultivars (EC_50_ = 0.88 – 3.42 mg/mL) (Nie et al. 2021). Their results further suggest potent nonartemisinin phytochemical(s) are present in both *Artemisia* species.

We and others noted there was anti-SARS-CoV-2 activity by other non-artemisinin antimalarial drugs including amodiaquine at an IC_50_ = 5.8 μM (this study) and 4.9-5.6 μM (Weston et al. 2020), tafenoquine at an IC_50_ of 2.6 μM (Dow et al. 2020), and lumefantrine at a reported IC_50_ = 23.2 μM (Cao et al. 2020). Gendrot et al. (2020) also reported anti-SARS-CoV-2 activity of various ACTs drugs at doses used for treating malaria with mefloquine-artesunate (550 mg + 250 mg, respectively) providing the maximum inhibition, namely 72% of viral replication at serum C_max_. Other combinations were less effective.

The high bioavailability of artemisinin after oral consumption of dried-leaf *A. annua* (DLA) was not surprising considering that a series of earlier studies in rodents showed the drug is >40 fold more bioavailable when delivered via the plant than in purified form (Weathers et al. 2011; Weathers et al. 2014). The increased bioavailability is mainly the result of three mechanisms: essential oils in the plant material improving the solubility of artemisinin, improved passage across the intestinal wall, and especially the inhibition of liver cytochrome P450s, 2B6, and 3A4 that are critical in first-pass metabolism (Desrosiers and Weathers 2016, 2018; Desrosiers et al. 2020). The anti-SARS-CoV-2 IC_90_ of the SAM1 and SAM2 cultivar samples ranged from 12.3-18.8 μM artemisinin, equal to 1.7-2.6 μg/mL, so 1 g of the SAM cultivar delivered *per os* yielded 2.6 μg/mL in a patient’s serum. Thus, 1 g of DLA could deliver enough artemisinin/DLA to achieve the IC_90_ of the hot water extract. While further human studies are required, this hypothetical estimation suggests that reasonable amounts of DLA consumed *per os* may deliver adequate amounts of the antiviral phytochemicals needed to provide a cost-effective anti-SARS-CoV-2 treatment. Indeed, the broad scale use of both artemisinin and non-artemisinin compound antimalarials including *A. annua* tea infusions across Africa may help in part explain why despite having anti-SARS-CoV-2 antibodies, Africans have not to date suffered the clinical scourge of SARS-CoV-2 like the rest of the world (Uyoga et al. 2020).

## 5.0 CONCLUSIONS

This is a first report of the *in vitro* activity against SARS-CoV-2 and two of its recent variants, BB1.1.7 and B1.351, by hot water extracts of *A. annua* at concentrations that can be achieved in humans following oral consumption of the plant material. Further work is required to isolate and identify the nonartemisinin bioactive phytochemicals in the plant extracts. If subsequent clinical trials are successful, *A. annua* could potentially serve as a safe therapeutic that could be provided globally at reasonable cost and offer an alternative to vaccines.

## Abbreviations

ACT: artemisinin combination therapy
ASAQ: artesunate amodiaquine
DLA: dried leaf Artemisia
DW: dry weight
EMEM: Essential Minimal Eagle’s Medium

## 6.0 ACKNOWLEDGEMENTS

We extend our gratitude to Tim Urekew (TJU Associates, NY, NY) and Scott Rudge (RMC Pharmaceutical Solutions, Inc., Longmont, CO), for their early advice and collaboration. Prof. David Ho is gratefully acknowledged for supporting the live virus work in his lab. We also thank Markus Hoffmann and Stefan Pöhlmann for providing the SARS-CoV-2 Spike Δ18 D614G plasmid. Award Number NIH-2R15AT008277-02 to PJW from the National Center for Complementary and Integrative Health funded phytochemical analyses of the plant material used in this study. The content is solely the responsibility of the authors and does not necessarily represent the official views of the National Center for Complementary and Integrative Health or the National Institutes of Health. SJP is partially supported by a Washington Research Foundation Technology Commercialization Phase 1 grant and NIH grant 3U41AT008718-07S1 from the National Center for Complementary and Integrative Health.

## 7.0 CONFLICT OF INTEREST STATEMENT

Authors declare they have no competing conflicts of interest in the study.

## 8.0 AUTHOR CONTRIBUTIONS

MSN – conducted SARS-CoV-2 experiments, helped analyze the data, and contributed to the manuscript.

YH – conducted SARS-CoV-2 experiments, helped analyze the data, and contributed to the manuscript.

DAF – provided reagents, helped analyze the data, and edited the manuscript. SJP – helped plan and analyze pseudovirus data, and contributed to manuscript

JW – conducted pseudovirus experiments, helped analyze data, and contributed to manuscript

MJT – prepared and analyzed plant extracts and human samples, helped analyze the data, and contributed to the manuscript.

PJW – wrote manuscript, conducted the single human PK test, provided reagents, and helped analyze the data.

## Supplemental Data

Bioavailability of artemisinin from per os consumption of dried leaf Artemisia annua in a human subject.

### NB

PJW verified with the WPI IRB that no IRB approval is required for self-experimentation.

### Method

One of the authors, PJW, age 71, 140 lb [63.6 kg]) consumed 3 g powdered, encapsulated dried *A. annua* SAM (2018 garden crop) and had 3 total blood draws: just prior to consumption; at 2 h post consumption, and a few weeks later, subject took another 3 g dose and blood was drawn 5 h post consumption. Serum was isolated from the blood and analyzed for artemisinin using gas chromatography mass spectrometry (GCMS) per Martini et al. 2020. Artemisinin (MW = 282.33) amount in the encapsulated material was 1.5% (15 mg/g), so amount consumed (delivered) was 45 mg artemisinin. Estimating 100% bioavailability, and that this human subject had a total volume of about 4.13 L blood (https://reference.medscape.com/calculator/estimated-blood-volume), the amount of delivered artemisinin/mL blood could not exceed 10.90 mg/L, or 10.90 μg/mL. Human blood is 55% serum (or 2.3 L for this human subject), so the highest serum concentration of artemisinin would actually be about 20 mg/L or 20 μg/mL.

**Table S1.** Human pharmacokinetics of ART delivered from *p.o. Artemisia annua*.^a^

|  |  |  |
| --- | --- | --- |
| Time (h) | ART in serum (µg/mL) | % of *A. annua*-ART consumed |
| 0 | 0.0 | 0 |
| 2 | 7.04 | 36 |
| 5 | 0.16 | 0.8 |

<sup>a</sup> Consumed 3 g powdered dried leaf *A. annua* containing 10.90 mg ART *in toto*; maximum possible serum concentration at *p.o.* delivery = 19.62 µg ART/mL.

**Table S2.** Examples of comparative amounts of artemisinin metabolites in various cultivars of *A. annua*.

|  |  |  |  |
| --- | --- | --- | --- |
| Compound | LUX<br>MNHNL17732 | BUR<br>LG0019527 | SAM1,<sup>1</sup><br>MASS 00317314 |
|  | mg/g dry weight leaves |  |  |
| Artemisinin | 1.34 | 1.70 | 10.94<sup>2</sup>, 15.90<sup>1</sup> |
| Arteannuin B | 0.93 | ND | 0.09<sup>2</sup>, 2.32<sup>1</sup> |
| Artemisinic acid | 0.86 | ND | ND<sup>2</sup>, 0.37<sup>1</sup> |
| Deoxyartemisinin | 0.32 | 0.39 | 0.83<sup>2</sup>, NM<sup>1</sup> |

ND, not detected; NM, not measured.
<sup>1</sup> Data from Weathers and Towler (2014); artemisinic acid is detectable in the dried leaves.
<sup>2</sup> Data estimated from fresh leaf analysis using dry weight/fresh weight ratio = 0.26 from Table 1 in Towler and Weathers (2015).

## Notes

### Competing Interest Statement

The authors have declared no competing interest.

### Summary of Updates

New data added to show efficacy against 2 new SARS-CoV-2 variants.

